# Insights into the catalytic properties of the mitochondrial rhomboid protease PARL

**DOI:** 10.1101/2020.07.27.224220

**Authors:** Laine Lysyk, Raelynn Brassard, Elena Arutyunova, Verena Siebert, Zhenze Jiang, Emmanuella Takyi, Melissa Morrison, Howard S. Young, Marius K. Lemberg, Anthony J. O’Donoghue, M. Joanne Lemieux

## Abstract

The rhomboid protease PARL is a critical regulator of mitochondrial homeostasis through its cleavage of substrates such as PINK1, PGAM5, and Smac, which have crucial roles in mitochondrial quality control and apoptosis. To gain insight into the catalytic properties of the PARL protease, we expressed human PARL in yeast and used FRET-based kinetic assays to measure proteolytic activity *in vitro*. We show PARL activity in detergent is enhanced by cardiolipin. Significantly higher turnover rates are observed for PARL reconstituted in proteoliposomes, with Smac being cleaved most rapidly at a rate of 1 min^−1^. PGAM5 is cleaved with the highest efficiency compared to PINK1 and Smac. In proteoliposomes, a truncated β-cleavage form of PARL is more active than the full-length enzyme for hydrolysis of PINK1, PGAM5 and Smac. Multiplex substrate profiling reveals a substrate preference for PARL with a bulky side chain Phe in P1, which is distinct from small side chain residues typically found with bacterial rhomboid proteases. This study using recombinant PARL provides fundamental insights into its catalytic activity and substrate preferences.

## Introduction

Mitochondria play an essential role in cellular respiration but also play an equally important role in modulating cell death(Vafai & Mootha, 2012). These functions rely on the selective quality control of mitochondrial protein homeostasis (proteostasis) (Baker et al, 2011), that includes the controlled turnover of regulators in the mitochondria by the mitochondrial intramembrane protease PARL. This enzyme was originally named <u>P</u>resenilin-<u>A</u>ssociated <u>R</u>homboid-<u>L</u>ike protease after discovery in a yeast-two hybrid screen (Chan & McQuibban, 2013; Pickrell & Youle, 2015). PARL cleaves various safeguards of mitochondrial health, including the kinase PINK1 (Phosphatase and tensin (PTEN)-induced putative kinase 1) (Jin et al, 2010; Meissner et al, 2011; Shi et al, 2011) and the phosphatase, PGAM5 (phosphoglycerate mutase family member 5) (Sekine et al, 2012`), both of which are known to play roles in mitophagy, the selective removal of damaged mitochondria (Lu et al). Hence, PARL has been renamed and the acronym now corresponds to <u>P</u>INK1/PGAM5 <u>A</u>ssociated <u>R</u>homboid-<u>L</u>ike protease (Spinazzi & De Strooper, 2016). Additional substrates of PARL have been identified in a recent proteomic analysis including the pro-apoptotic factor Smac (Saita et al, 2017). The PARL knockout (KO) mouse results in a severe respiratory defect, similar to Leigh’s syndrome, which is a consequence of misprocessing of the nuclear encoded substrate TTC19, a subunit of complex III (Spinazzi et al, 2019). These PARL KO mice present with a severe motor defect, the loss of grey matter (cell bodies of neurons) in the cortex, and early lethality. The mitochondria of the KO mice have a distinct morphology lacking cristae that precedes neurodegeneration in grey matter(Cipolat et al, 2006; Frezza et al, 2006). When the PARL orthologue Pcp1/Rbd1 is knocked out in yeast, similar disturbances are also observed where the cristae and protein-mtDNA assemblies in the matrix dissipate (Herlan et al, 2003; McQuibban et al, 2003). These studies emphasize the essential nature of the PARL-type proteases for cell viability across evolution.

The PARL protease is a member of the rhomboid intramembrane protease family, which are membrane-embedded serine peptidases that are thought to be constitutively active. Their functions range from cleavage and release of membrane-tethered signaling molecules to membrane protein degradation (Kuhnle et al, 2019; Ticha et al, 2018). Regulation of PARL activity at the molecular level is thought to occur via post-translational modifications. Different forms of PARL have been identified in various tissues as a result of processing events; these include a protein with a mitochondrial matrix targeting sequence (MTS), a full-length mature form after removal of the MTS (PARLΔ55) and a truncated form derived from cleavage at residue S77 (PARLΔ77), referred to as β-cleavage (Jeyaraju et al, 2011). Ectopic expression of this truncated form of PARL in tissue culture was shown to alter mitochondrial morphology leading to fusion defects and hence has been suggested to be more active(Jeyaraju et al, 2011). In contrast, it was shown by others that truncation of PARL lead to decreased processing of PINK1 (Shi et al, 2011). The truncation site at S77 was shown to be phosphorylated in response to stress, an event that influences β-cleavage of PARL and its activity in tissue culture cells (Shi & McQuibban, 2017). The mechanism of this putative regulatory switch remains to be determined and it has not yet been established if S77 PARL is generated by PARL itself or other mitochondrial proteases. Despite this, PARL has not been characterized at the molecular level and thus the importance of β-cleavage in its regulation remains unclear.

Our analysis with human recombinant PARL, comparing full-length and the truncated β-cleavage form, allows us to determine kinetics of substrate cleavage and examine the parameters influencing PARL activity. We evaluated PARL activity using SDS-PAGE and cleavage of peptide substrates using FRET-based fluorescent assays. We observe that β-cleavage increases the catalytic activity of PARL. When PARL is reconstituted in a lipid environment similar to that found in the inner mitochondrial membrane (IMM), we reveal similar substrate specificities yet an enhanced catalytic rate of cleavage of all substrate peptides when compared to those measured with the enzyme in detergent micelles. In addition, we observe that PARL activity was highly increased by cardiolipin (CL). Together this work provides characterization of the PARL protease and further extends our mechanistic understanding of this important safeguard of mitochondrial homeostasis.

## Results

### Recombinant human PARL expressed in yeast is active

To examine the molecular features that determine PARL activity, we took an approach to express and purify recombinant PARL and study it *in vitro*. PARL is a nuclear-encoded serine protease with an N-terminal MTS. Human PARL was cloned into a His-tagged expression vector where we added a C-terminal GFP-tag, which allowed us to determine conditions for its optimal expression in *Pichia pastoris*. Since the PARL precursor with the MTS intact resulted in poor yield (results not shown), both the full-length mature form (PARLΔ55) and a truncated version representing β-cleavage (PARLΔ77) were expressed. (**Fig 1A, Supplemental Fig1,2**). In addition, the active site mutant PARL-Δ77-S277A was also generated. Recombinant PARL proteins were purified using affinity chromatography from dodecylmaltoside (DDM)-solubilized membrane fractions, a mild detergent that is known to result in low delipidation during extraction (Lemieux et al, 2002), followed by removal of the GFP-His-tag. Milligram quantities of all PARL variants were obtained using this expression system.

**Figure 1.**
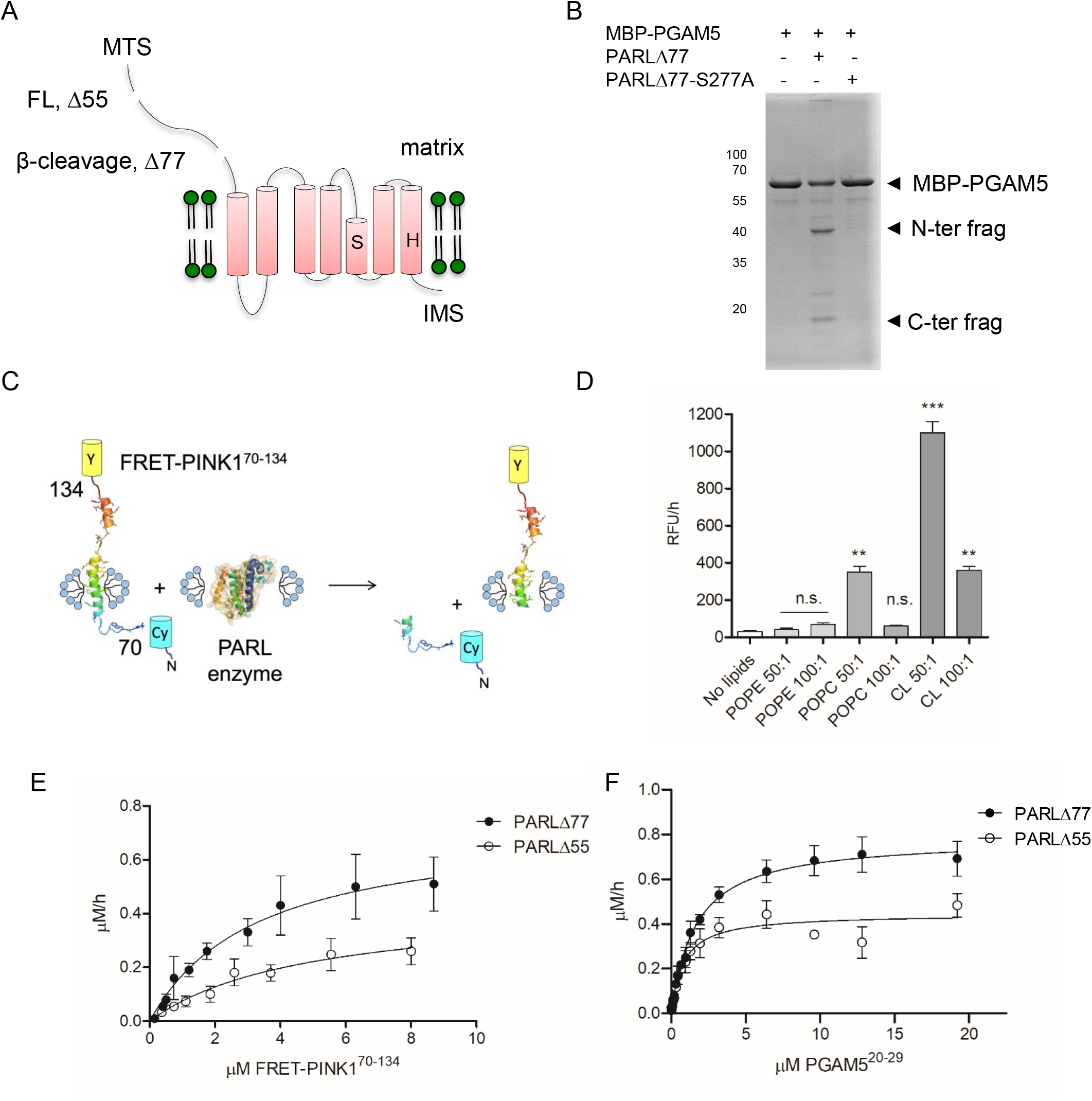
Recombinant human PARL protease is active. **A.** Cartoon representation of the PARL protease topology and truncations. Recombinant human PARL was expressed in *P. pastoris* to generate full-length (FL) starting at residue 55 or the β-cleavage form truncated at residue 77. An inactive PARL-Δ77-S277A mutation was also generated. **B.** Incubation of recombinant PARL with MBP-PGAM5 reveals an expected shift on SDS-PAGE **C.** A cartoon representation of the substrate construct with residues 70-134 of PINK1 flanked by the CyPet/YPet fluorescence reporter pair. **D.** Cleavage of FRET-PINK1(70-134) by detergent solubilized PARLΔ77 in the presence of increasing lipids. N=3 * p < 0.05; ** p < 0.005, *** p < 0.0005 n.s. denotes no significance **E.** Representative Michaelis-Menten curves for FRET-PINK170-134 cleavage by PARLΔ77 and PARLΔ55 N=3. **F.** Representative Michaelis-Menten kinetic curves for IQ-PGAM5 substrate cleavage by PARLΔ77 and PARLΔ55. N=3. For all experiments, values are represented as mean ±SEM.

To assess if the recombinant PARL was active, we first examined the cleavage of the transmembrane (TM) domain of PGAM5 fused to an N-terminal maltose binding protein (MBP) and a C-terminal Thioredoxin 1 domain (**Fig 1B**). The approach of using a substrate TM segment fused to MBP was undertaken before for both eukaryotic and prokaryotic rhomboids and the presence of the tag did not affect their ability to cleave the substrates (Strisovsky et al, 2009; Ticha et al, 2017a; Torres-Arancivia et al, 2010). Upon incubation of the MBP-PGAM5 fusion protein with PARLΔ77 in presence of CL to increase rhomboid activity (see below), new bands, representing the N- and C-terminal cleavage products, were observed on SDS-PAGE. These bands are not present following incubation of MBP-PGAM5 with the inactive S277A protein. This assay confirms the functionality of human mitochondrial PARL generated in the *P. pastoris* system.

### Lipids enhance PARL activity

To assess the catalytic properties of PARL protease and factors influencing its activity, we designed a FRET-PINK1^70-134^ substrate with fluorophores that were previously used to assess bacterial rhomboid protease activity *in vitro*(Arutyunova et al, 2014; Arutyunova et al, 2016). Residues 70-134 of PINK1, encompassing the predicted TM segment (residues 89-111) and adjacent residues, were cloned between two fluorescent protein reporters, YPet and CyPet, to allow for FRET activity measurement upon cleavage (**Fig 1C**) (Alford et al, 2013).

Lipids are known to influence membrane protein function(Bondar & Lemieux, 2019). In mitochondria, for example, increased CL amounts are observed during mitochondrial stress which influences protein function at the molecular level (Ruggiero et al, 1992). Therefore, we assessed the ability of β-cleavage PARL (PARLΔ77) to process the FRET-PINK^70-134^ substrate in the presence of the three primary lipids present in the IMM, namely CL, phosphatidylcholine (POPC), and phosphatidylethanolamine (POPE) (**Fig 1D)**(Comte et al, 1976). When compared to conditions with no lipid added, POPE at a molar ratio of 50:1 and 100:1 did not significantly increase PARL activity while POPC at a molar ratio of 50:1 increased activity by 4-fold. In addition, CL at a molar ratio of 50:1 caused a 100-fold increase of PINK1 cleavage, while only a 4-fold increase at a molar ratio of 100:1 (**Fig 1D).** This suggests that CL may modulate the activity of the mitochondrial rhomboid protease PARL by enhancing its overall structural stability through protein-lipid interactions, similar to other mitochondrial proteins (Ghosh et al, 2020; Ryabichko et al, 2020). It is also worth mentioning that CL is known to induce curvature in membranes (McMahon & Boucrot, 2015), which could alter the activity of PARL protease. Therefore, CL was included in all subsequent protein preparations of detergent solubilized PARL.

The cleavage of FRET-PINK1^70-134^ by DDM-solubilized PARLΔ55 and PARLΔ77 in the presence of CL obeyed Michaelis-Menten kinetics (**Fig 1E, Supplemental Table 1),** and revealed slow rates of cleavage, 0.46±0.09 h^−1^ and 0.43±0.06 h^−1^ respectively. This however reflects the tendency of intramembrane proteases to have slow turnover rates (Arutyunova et al, 2014; Kamp et al, 2015).

To further evaluate the cleavage of other known substrates of PARL, we generated internally quenched (IQ) peptide substrates (Arutyunova et al, 2018; Lapek et al, 2019; Naing et al, 2018; Ticha et al, 2017a) based on the amino acids flanking the PARL cleavage sites of PINK1(Deas et al, 2011), PGAM5 (Sekine et al, 2012), and Smac (Saita et al, 2017). Kinetic analysis using both full-length and β-truncated PARL with IQ-PINK1^99-108^, IQ-PGAM5^20-29^, and IQ-Smac^51-60^ peptide substrates was first performed in detergent, which revealed similar Michaelis-Menten kinetics for all peptides (**Fig 1F**). The assay allowed us to determine the catalytic parameters for the three primary PARL substrates and examine the substrate specificity (**Fig 2**, **Supplemental Tables 2,3 and 4)**. As shown above, a slow turnover rate for all substrates was observed, with the k_cat_ values ranging from 0.5 to 1.3 h^−1^ with the Smac peptide being the fastest cleaved and PGAM5 being the most efficiently cleaved based on k_cat_/K_M_ value.

**Figure 2.**
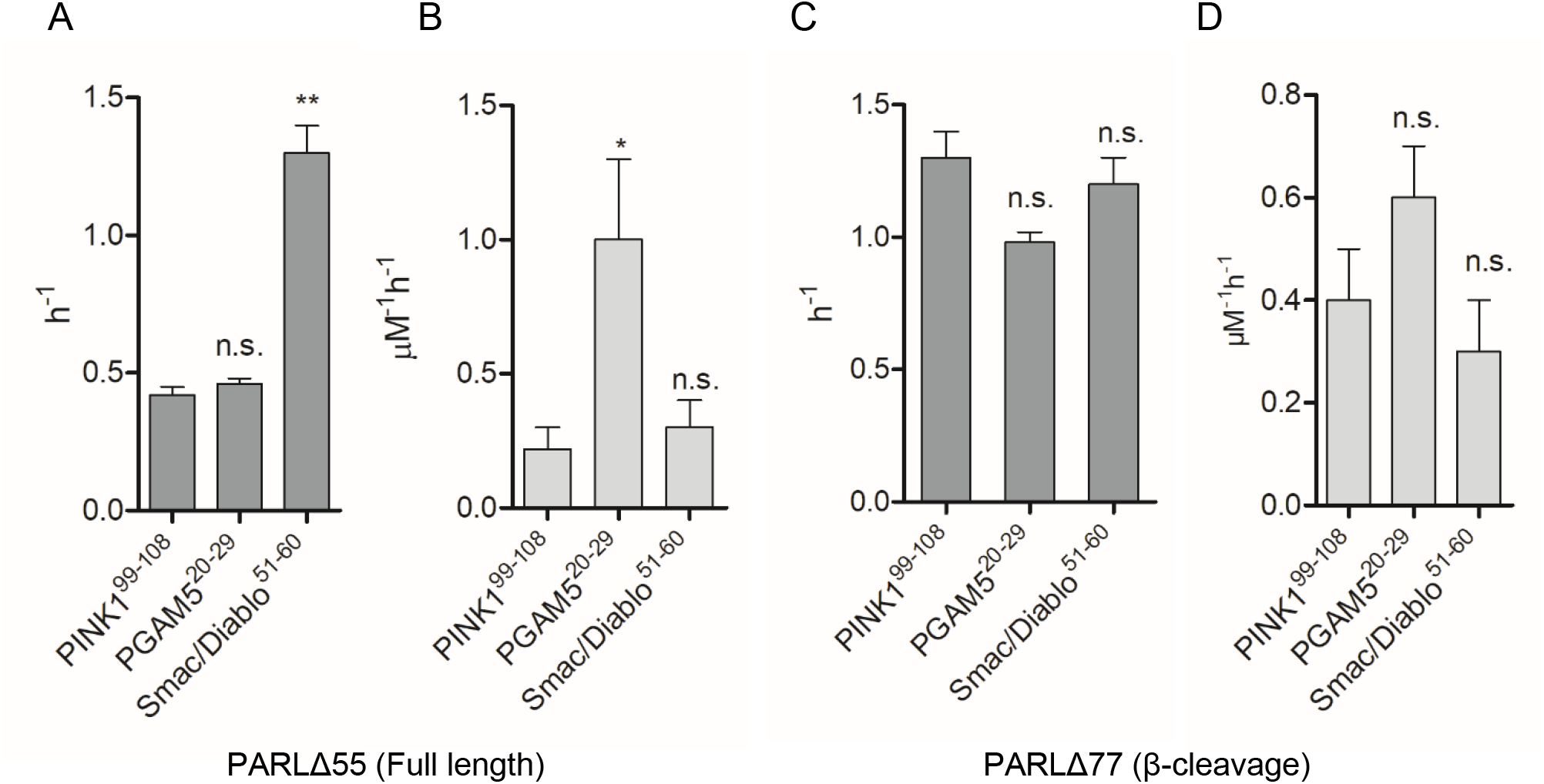
Catalytic parameters for IQ-peptide substrate cleavage with both PARLΔ55 (full length) and PARLΔ77 (β-cleavage) reveal differences in k_cat_ and k_cat_/K_M_ values. **A.** turnover rates and **B.** catalytic efficienty are plotted in a bar graph for full-length PARL. **C.** turnover rates and **D.** catalytic efficienty are plotted in a bar graph for β-cleaved PARL. Experiments were conducted in duplicate with an N=3-5. Values are represented as mean ±SEM (* p < 0.05; ** p < 0.005, n.s. denotes no significance).

These data also revealed that the turnover rates of the β-truncated PARL protease, with the short IQ-PINK1^99-108^ peptide, (0.42±0.03 h^−1^), and longer FRET-PINK1^70-134^ substrate (0.46±0.09 h^−1^) (**Fig 1E, Supplemental Tables 1 and 2)** are comparable. We conclude that the regions adjacent to the PINK1 cleavage site do not influence the cleavage process, and that the IQ-peptide substrates are suitable for performing further kinetic studies.

### PARL shows enhanced catalytic rate in liposomes

To assess the activity of recombinant PARL towards known substrates in a lipid bilayer, full-length and β-truncated PARL were reconstituted in proteoliposomes using *E. coli* lipids that closely resemble the composition of the IMM(Comte et al, 1976). To calculate the specific activity of the protease we quantified the fraction of PARL with an outward facing active site using an activity-based TAMRA-labeled fluorophosphonate probe (Sherratt et al, 2012). We revealed that ~70% of PARL was oriented in a substrate-accessible, outward-facing manner (**Supplemental Fig 3**). Next, we compared cleavage of our three IQ-peptide substrates by reconstituted full-length mature PARLΔ55 and the β-truncated form, PARLΔ77 (**Fig 3, Supplemental Tables 5 and 6).** Overall, we observed that the lipid environment increased the activity of both forms of PARL towards all substrates and the reaction still displayed Michaelis-Menten kinetics (**Fig 3C and Supplemental Table 5 and 6)** with only negligible background activity detected for PARLΔ77-S277A (**Supplemental Fig 4**). Of all peptides assessed, cleavage of Smac by β-truncated PARL#x0394;77 was the fastest with a turnover rate of 58 ±7 h^−1^, or 1.0±0.1 min^−1^, and PGAM5, was the preferred substrate with the lowest K_M_ and the highest catalytic efficiency (k_cat_/K_M_) of 26 ± 8 μM^−1^h^−1^ (**Supplemental Table 6**), which is in agreement with the data obtained in a detergent environment.

**Figure 3.**
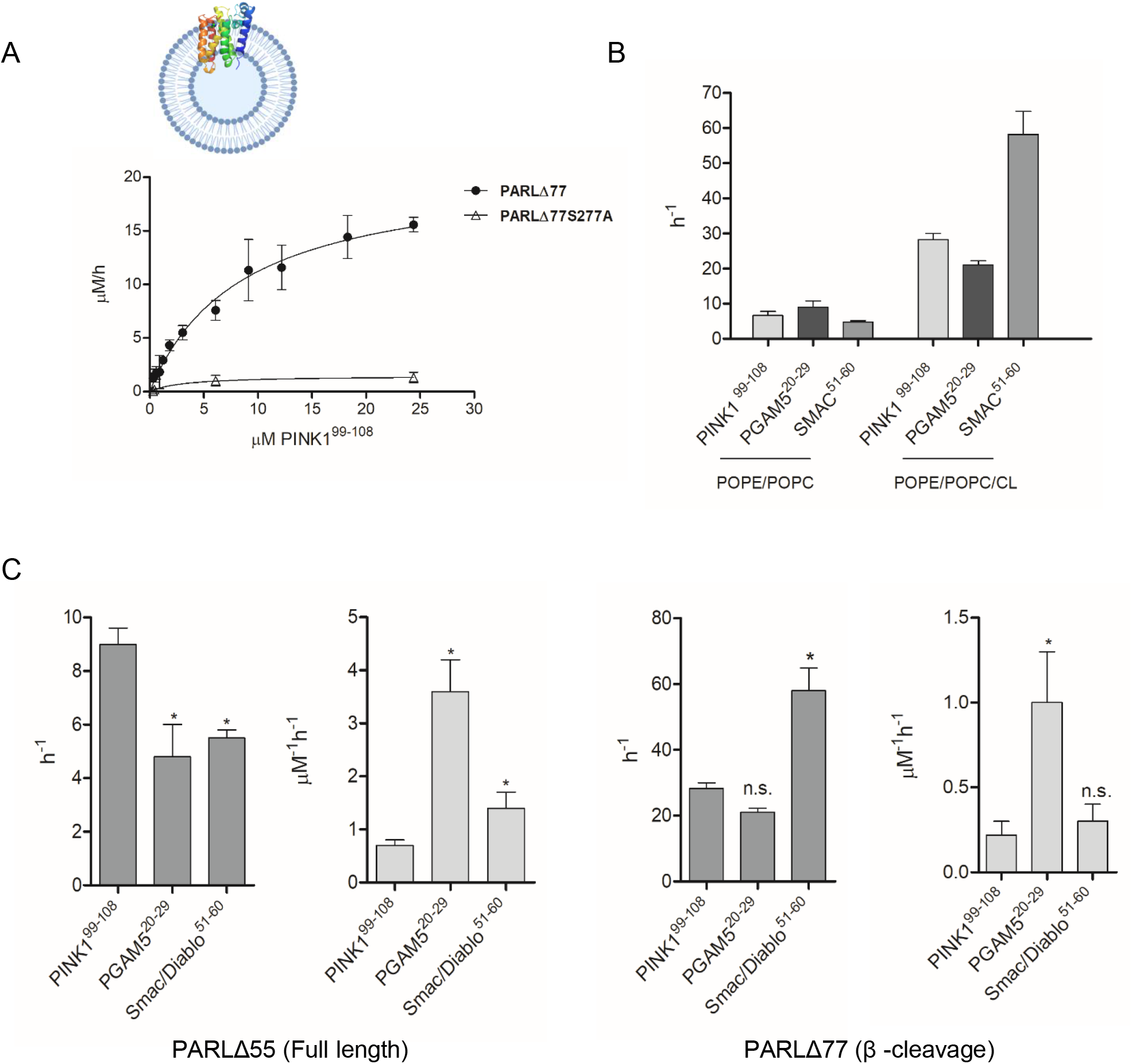
Enhanced catalytic rate is observed with PARL in proteoliposomes with cardiolipin. **A.** Representative Michaelis-Menten curve for PARLΔ77 cleavage of IQ-PINK199-108. **B.** The effect of CL on PARLΔ77 activity in proteoliposomes with IQ-peptide substrates. **C.** Bar graphs for turnover rates and catalytic efficiencies of proteoliposome-reconstituted PARLΔ55 and PARLΔ77 with IQ-peptide substrates. Experiments were conducted in duplicate with an N=3-5. Data is represented as mean ± SEM (* p < 0.05; ** p < 0.005, n.s. denotes no significance).

**Figure 4.**
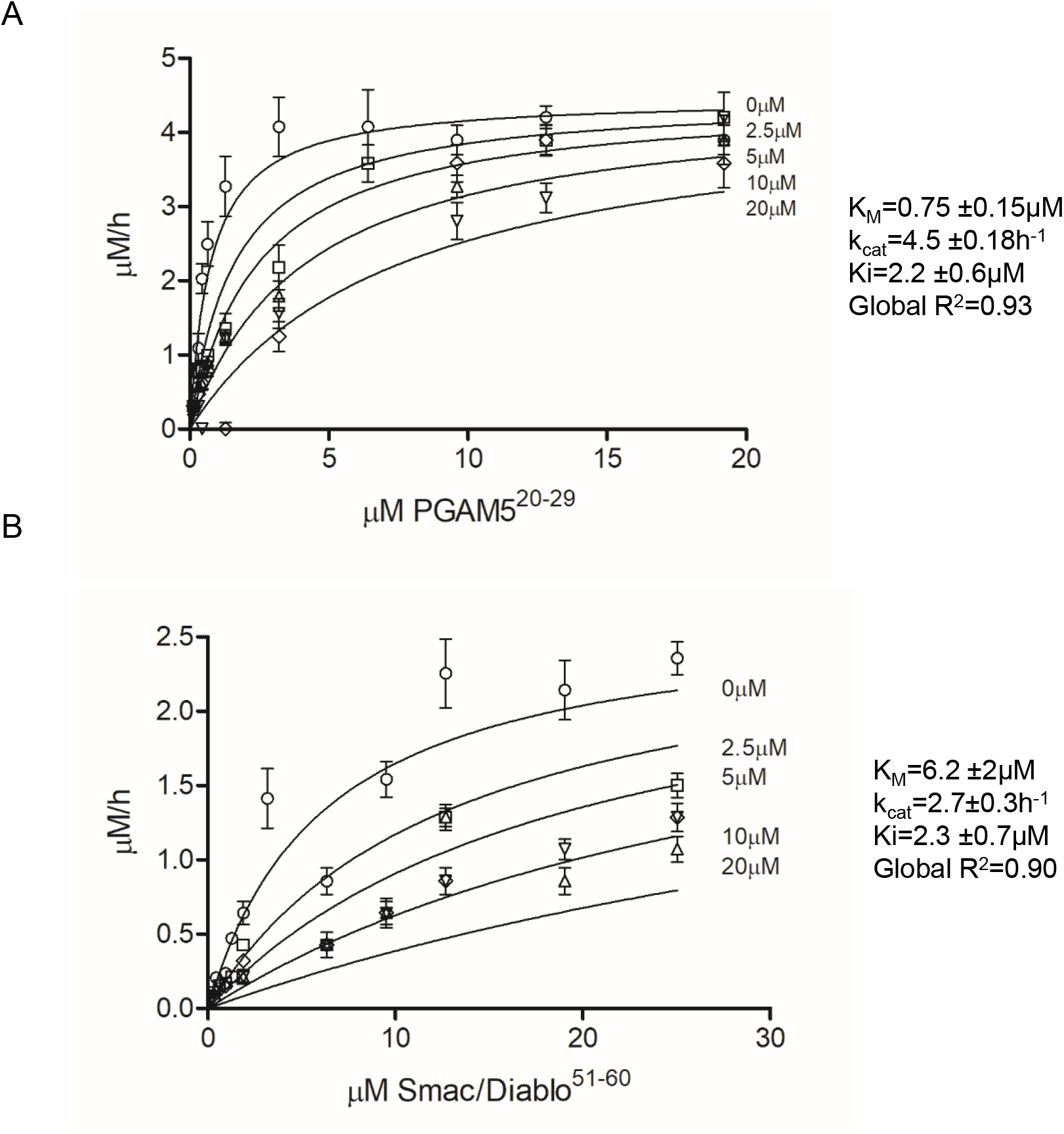
Competitive studies of PARL77-mediated cleavage of PGAM5^20-29^ and SMAC^51-60^ peptides in the presence of PINK1^89-111^ reveals competitive inhibition. Cleavage assays of PARL77 (1μM) with: A. IQ -PGAM5^20-29^ (0.13-19 μM) B. or SMAC^51-60^ (0.3-25μM), performed in the presence of different concentrations of non-fluorescent PINK1^89-111^ substrate (2.5, 5,10, 20μM), reveals competitive inhibition suggestive of identical binding sites. Fluorescence detection of each substrate concentration in the presence of corresponding PINK1^89-111^ without enzyme was used as a negative control. Initial velocities were determined for each substrate concentration. Michaelis–Menten plots were subjected to global fit to distinguish the kinetic model and determine the kinetic parameters. Values are represented as mean ±SEM.

To confirm the effect of CL on PARL activity in proteoliposomes we prepared two types of proteoliposome samples containing β-truncated PARL and conducted detailed kinetic analysis with the three IQ-peptide substrates (**Fig 3B**). The first sample consisted only of POPC and POPE, the second type also contained CL. The turnover rates for the PINK1, PGAM5 and Smac peptide substrates with the two PL samples demonstrated a trend similar to that observed in the detergent environment (**Fig 3C)**. When cardiolipin was omitted from the liposomes, we observed 2- to 10-fold slower turnover rates of cleavage for our three IQ-peptide substrates by β-truncated PARL (**Fig 3B, and Supplemental Table 7**). This result again demonstrates that CL, which is specific to the inner mitochondrial membrane where PARL resides have an effect on the proteolytic activity of PARL.

It should be noted that the IMM is the only eukaryotic cellular membrane that contains a significant amount of CL, which is known to be essential to the activity of numerous IMM proteins. There are currently 62 different proteins reported to interact with CL and high-resolution structures of with at least one CL molecule present (Planas-Iglesias et al, 2015) CL is a structurally unique lipid composed of four acyl chains and two phosphates, and both hydrophobic and polar groups can both potentially hold a negative charge it can form lipid-protein interactions through phosphate binding, hydroxyl binding, or acyl binding patches on a membrane protein (Musatov & Sedlak, 2017) Whether specific lipid-protein interactions result in increased stability of PARL or an actual enhancement of its function, or perhaps a combination of both, remains to be elucidated.

### β-cleavage influences the activity of PARL

Our detailed kinetic analysis also allowed us to examine whether β-cleavage influences the activity of PARL *in vitro*. We have determined kinetic parameters of full length and β-truncated PARL in both detergent (**Fig 2, Supplemental Tables 3 and 4**) and proteoliposomes (**Fig 3, Supplemental Tables 5 and 6**), using three different IQ-peptide substrates: PINK1, PGAM5, and Smac. In detergent, turnover rates increase (PINK1 and PGAM5) or are unchanged (Smac) with β-truncated PARL when compared to full length-PARL (**Fig 2**). In proteoliposomes, with bilayer environment, a similar trend is observed for PINK1 and PGAM5, with the turnover rate of Smac also being enhanced 10-fold with β-truncated PARL (**Fig 3 and Supplemental Table 8**). Overall this suggests β-cleavage, i.e. truncation at S77, results in an enhancement of substrate cleavage, however there seem to be substrate-dependent effects.

The influence of PARL truncations on activity has been controversial. Processing of PARL to either its full-length form, predicted to occur at Δ53, or the further β-truncated PARLΔ77 form, has been proposed to be a modulator of its enzymatic activity (Sik et al, 2004). Cellular studies have shown impaired PARL cleavage of PINK1 when a mutation at Ser77 is introduced that prevents truncation to the PARLΔ77 form (Shi et al, 2011; Shi & McQuibban, 2017), with additional data suggesting the longer form is less active towards PINK1. Another study showed that the β-truncated PARL (PARLΔ77) induces mitochondria fragmentation in cells (Jeyaraju et al, 2011). We observe higher activity with the shorter β-cleavage form of PARL, PARLΔ77. However, our data reveals that compared to full length, in detergent truncated PARL results in only a 2-3-fold enhancement of substrate cleavage for PINK1 and Smac, and no change is observed with the PGAM5 substrate. A 10-fold enhancement with Smac in proteoliposomes is observed (**Supplemental Table 8**). The direct influence of PARL truncations on substrate cleavage has not been examined *in vitro* before and with our data we confirm that PARL is catalytically active in either form, thus indicating that processing to the β-cleavage form is not required for proteolytic activity or PARL functionality as was once speculated(Shi et al, 2011).

### PARL is weakly inhibited by commercial inhibitors

Rhomboids were initially discovered to be serine proteases because the first identified rhomboid protease from *Drosophila* Rhomboid-1 was sensitive to serine protease inhibitors dichloroisocoumarin (DCI) and tosyl phenylalanyl chloromethylketone (TPCK). Further, the crystal structures of bacterial rhomboids with serine protease inhibitors – diisopropylfluorophosphate, isocoumarins aided in revealing structural insight about molecular mechanism of catalytic reaction (Ticha et al, 2018). However, the broad-spectrum serine protease inhibitor PMSF does not act on rhomboid proteases (Lemberg et al; Urban et al; Urban & Wolfe, 2005). Inhibition of PARL has not been previously explored.

We examined three standard serine protease inhibitors families: sulfonyl fluoride (PMSF), chloromethyl ketone (TPCK), and coumarin (DCI), with our *in vitro* PARL assay. In proteoliposomes, for both PINK1 and PGAM5 peptide substrates, we show that PARL activity is not inhibited by 100 μM PMSF, whereas 100 μM TPCK partially inhibits and 100 μM DCI has the largest inhibitory effect (**Supplemental Fig 6**), which is in agreement with previously shown effect for bacterial rhomboid proteases (Strisovsky, 2016). Overall this supports the view that for PARL, specific inhibitors will need to be developed similar to bacterial rhomboid proteases (Ticha et al, 2017b; Verhelst, 2017).

### PINK1, PGAM5 and Smac are competing for the same binding site

With multiple substrates discovered, it is obvious that PARL has pleotropic roles in mitochondria and that substrate cleavage must be precisely regulated(Lysyk et al, 2020), whether though compartmentalization(Wai et al, 2016), or differential substrate binding. It was shown that PARL mediated differential cleavage of PINK1 and PGAM5 depends on the health status of mitochondria (Wai et al, 2016). Further studies demonstrated that the rate of PINK1 cleavage in cells is influenced by PGAM5, indicating that PINK1 and PGAM5 may compete for cleavage by PARL(Lu et al, 2014). However, it has been speculated that PINK1 and PGAM5 are not competitive substrates *in vivo* since reducing the expression of PINK1 by siRNA did not increase cleavage of PGAM5 by PARL (Sekine et al, 2012), highlighting that function of PARL changes in response to ΔΨ_m_ loss. This raises questions whether PARL substrates bind to the same residues in the active site or alternative binding sites might exist on enzyme’s surface.

To address this, we performed competition binding assays using fluorescent IQ-PGAM5^20-29^ and IQ-Smac^51-60^ as the main substrates and non-fluorescent PINK1^89-111^ (^89^ AWGCAGPCGRAVFLAFGLGLGLI^111^) as a competing substrate (**Fig 4**). The Michaelis-Menten curves of PARL-mediated cleavage of IQ-PGAM5^20-29^ and IQ-Smac^51-60^ were obtained in the presence of different concentrations of PINK1^89-111^. The data sets were fitted globally to competitive, non-competitive, and mixed inhibition with strong preference for competitive inhibition for both substrates and global R^2^ of 0.93 and 0.90 for IQ-PGAM5^20-29^ and IQ-Smac^51-60^ respectively. The determined K_d_ for IQ-PINK1^89-111^ (represented by K_i_ of the PARL-PINK^89-111^ complex) was 2.4 ± 0.6 μM when IQ-PGAM5^20-29^ was used as the main substrate and 2.3 ± 0.7 μM when IQ-Smac^51-60^ was used (Fersht, 2002). These competitive inhibition parameters between the two substrates revealed that the assessed PARL substrates bind to the same binding site with similar affinities and are exclusive to each other. These results suggest that the inverse regulation of PINK1, PGAM5 and Smac cleavage observed in cells is controlled by other mechanisms like compartmentalization, involvement of protein partners for substrate presentation or different accessibility of scissile bond in response to different membrane conditions.

### PARL has bulky substrate specificity preferences distinct from bacterial rhomboid proteases

Bacterial rhomboid proteases are known to cleave substrates with some specificity for small side chain residues in the P1 position and bulky hydrophobic residues in the P4 position(Strisovsky et al, 2009). Analysis of the C-terminal cleavage products of PARL-catalyzed cleavage isolated from cell extracts by Edman degradation (Deas, 2011; Sekine 2012) or mass spectrometry (Saita, 2017) so far revealed no consensus. Using recombinant PARL, we now assessed the cleavage site specificity using a library of 228 synthetic peptides that are each 14 amino acids in length. We have previously confirmed that this library contained many peptide substrates that are cleaved by the bacterial rhomboids from *Providencia stuartii* and *Haemophilus influenza* (Lapek et al, 2019). PARL in buffer containing either DDM or reconstituted in proteoliposomes was incubated with an equimolar mixture of peptides and 70 common cleavage products by the enzyme under these two different assay conditions were identified (**Fig 5**). These data indicated that the environment for PARL may alter the substrate cleavage preference of the enzyme or prevent access of certain substrates to the active site. We analyzed the location of cleavage within the peptide substrates and discovered that when PARL is assayed in proteoliposomes, it cleaves many peptides near the amino terminus, while PARL in DDM does not (**Fig 5B**). Importantly, analysis of the amino acids surrounding the cleaved bond shows that PARL when assayed in either DDM or proteoliposomes, no small amino acid is found in the P1 position. In DDM PARL preferentially cleaves peptides with norleucine (Nle), Tyr, Phe, Arg, and Lys in the P1 position while Phe and Ile are most frequently found in the P1ʹ position (**Fig 5A**). In addition, hydrophobic amino acids are found at the P4 and P2 position. When PARL is assayed in proteoliposomes, Phe and Nle are found most frequently in the P1 and P1ʹ positions (**Fig 5B**). Lys is also significantly enriched in the P1 position, while hydrophobic amino acids are found in the P4, P2, and P4ʹ positions. When these profiles are compared to the sequences surrounding the putative PINK1 cleavage site, VFLA-FGLG only P1ʹ-Phe and P4-Val are well tolerated from this sequence. In fact, Ala at P1 and Gly at P2ʹ are disfavored by PARL when assayed in DDM and proteoliposomes, respectively.

**Figure 5.**
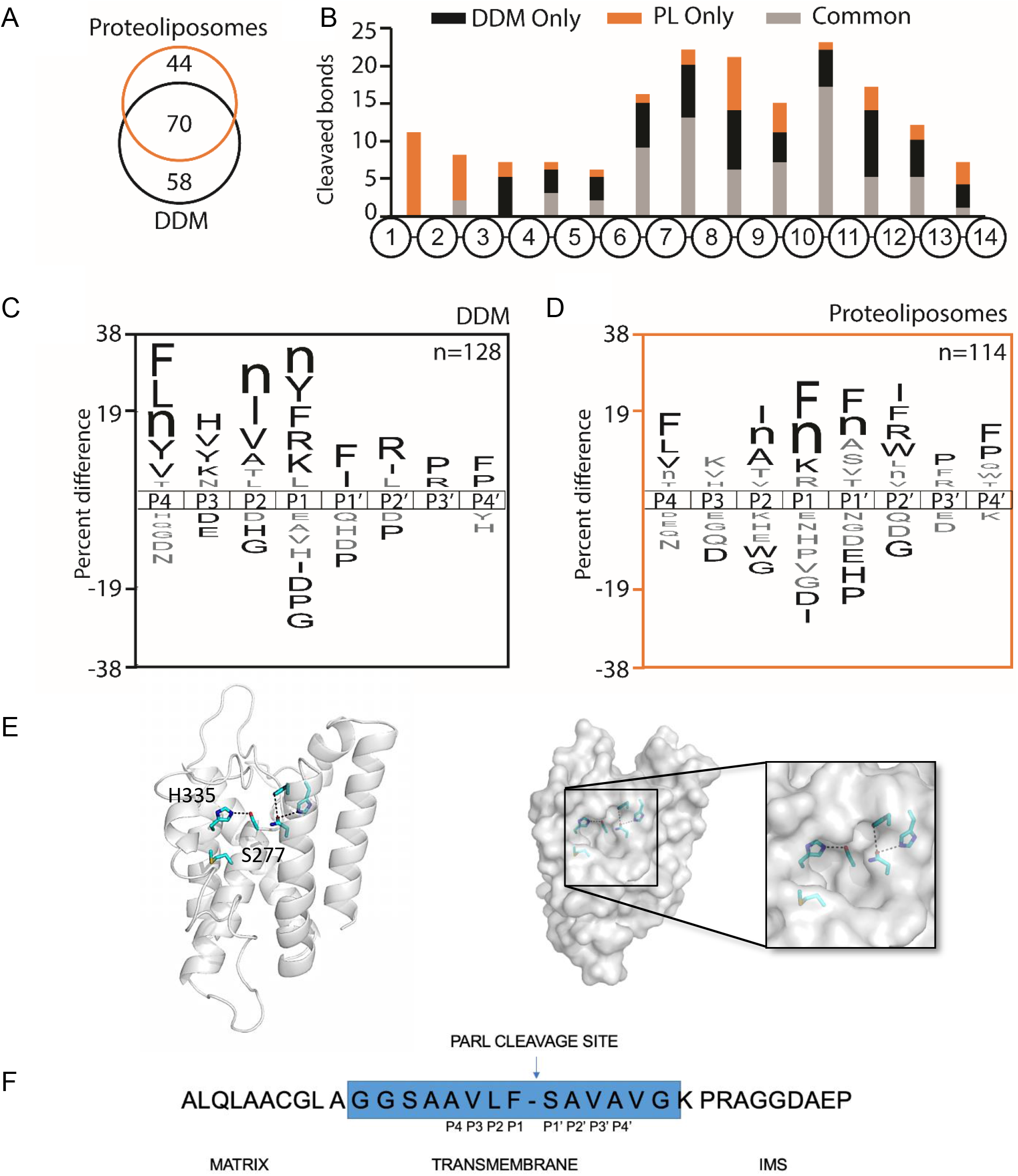
PARL protease has substrate specificity distinct from bacterial rhomboid proteases. Substrate specificity parameters using multiplex LC-MS/MS screening with 283 substrates. Samples assessed were PARL in DDM with CL added, and PARL in PL containing PC, PE and CL. **A**. Common peptides cleaved between PARL in DDM (Black line) and PL-reconstituted PARL (Orange line) **B**. Position of cleavage of IQ-peptide between DDM and PL-PARL **C.** Amino acid preferences for PARL in DDM vs **D.** Parl in PLs. **E.** PARL homology model based on the HiGlpG structure (2NR9.pdb) reveals a helical bundle and catalytic Ser277-His355 dyad. A surface representation reveals a P1 pocket near the catalytic serine. **F.** N-terminal sequencing of the C-terminal MBP-PGAM5 cleavage product reveals a Phe residue in the P1 position. Blue represents the predicted TM boundary of PGAM5.

A GlpG-based homology model of human PARL reveals a putative pocket in the P1 position that could accommodate such a bulky side chain (**Fig 5C**). The P4 position has a consistent bulky residue similar to that for bacterial rhomboid proteases. When PARL was incubated with the full TM region of MBP-PGAM5, N-terminal sequencing revealed that cleavage occurred between Phe-23 and Ser-24 (**Fig 5D**). Interestingly, this cleavage site determined in our *in vitro* assay is different from the sites previously determined in tissue culture cells, which for PGAM5 was cleaved between Ser-24 and Ala-25 (Sekine et al, 2012). As the readout of an *in vitro* assay is more direct as determination of N-terminus of cleavage fragments isolated from tissue culture cells, we suggest the *in vivo* cleavage fragments may become subject to further trimming by additional proteases. However, our *in vitro* assay revealed that the PGAM5 cleavage site, originally thought to be in the center of the TM domain, topologically, are now placed closer to the matrix-exposed rhomboid active site. Taken together with the results from the peptide library (**Fig 5A andB**), the *in vitro* determined PGAM5 cleavage sites support the preference for PARL to cleave proteins and peptides with a bulky amino acid such Phe in the P1 position.

This is the first substrate specificity study of PARL which shows an interesting preference for Phe in the P1 position. For bacterial rhomboid proteases, small side chain residues are preferred in the P1 position(Urban & Freeman, 2003). In fact, no cleavage of the TatA substrate occurs by bacterial rhomboid protease AarA when Phe is mutated into the P1 position of the substrate cleavage site (Strisovsky et al, 2009). Analysis of the *E. coli* rhomboid protease with a peptide substrate transition analog revealed a similar preference (Zoll et al, 2014). Thus far, YqgP from *Bacillus subtilis* is the only bacterial rhomboid protease known to cleave with Phe at the P1 position (Strisovsky et al, 2009). YqgP is evolutionarily distinct from the *E. coli* GlpG {Lemberg, 2007 #149}, which suggests evolutionary pressure on substrate specificity. With many mitochondrial proteins harboring Phe residues in their TM regions, PARL may have co-evolved a different mode of substrate recognition. This is the first substrate preference determined for any eukaryotic rhomboid protease and the preferences for others, such as the Golgi RHBDL1 and ER RHBDL4 (Kuhnle et al, 2019), remain to be determined.

## Materials and methods

### Expression and purification of recombinant PARL

PARL gene (PARLΔ77, PARLΔ77-S277A, or PARLΔ55) was cloned into pPICZA-GFP vector with a C-terminal hexahistidine-tag (Brooks et al, 2013). An identified high-expressing clone was grown overnight at 28°C in 100 mL of BMGY media to an OD_600_ of 4. A total of 6 L of BMGY media was sub-inoculated to a starting OD_600_ of 0.03 and grown for 20 h at 28°C. Cells were harvested by centrifugation and cell pellets were resuspended in an equal volume of BMMY induction media. Cultures were induced for 48 h at 24°C, with fresh methanol being added after 24 h to a final concentration of 1% (v/v). Cells were harvested and resuspended in TBS buffer (50 mM Tris-HCl pH 8.0, 150 mM NaCl). Cells were re-suspended in TBS with PMSF and lysed by passage through a Constant Systems cell disruptor at 38.2 kPSI, and membranes were isolated by ultracentrifugation. Membranes were homogenized in 50 mM Tris-HCl pH 8.0, 200 mM NaCl, 5% glycerol, 20 mM imidazole, 1 mM PMSF, and solubilized using 1.2% Triton X-100. Insoluble material was pelleted by ultracentrifugation and the supernatant bound to HisPur™ cobalt resin (ThermoFisher, USA) by gravity flow-through column. The protein-bound resin was washed with 10 mM imidazole and eluted with imidazole (50 mM Tris-HCl pH 8.0, 300 mM NaCl, 20 % glycerol, 0.1% DDM (Anatrace, USA), 1 M imidazole). The purified PARL-GFP fusion protein was digested by incubation with TEV protease and 1 mM TCEP overnight at 4°C. Dialysis was performed for 2 h to remove imidazole and TCEP (50 mM Tris-HCl pH 8.0, 300 mM NaCl, 20% glycerol). PARL was purified from GFP and TEV using HisPur™ Ni-NTA agarose resin (ThermoFisher, USA). Flow-through was collected and concentrated using a 10 000 MWCO concentrator (Millipore, USA). Protein concentration was determined by BCA assay (Pierce™ BCA Protein Assay Kit, ThermoFisher, USA). Purified protein was incubated on ice with dried cardiolipin (Sigma-Aldrich, USA) to a final lipid concentration of 0.1 mg/mL. Protein-lipid sample was aliquoted, flash frozen, and stored at −80°C.

### FRET-PINK1 purification

Residues 70-134 of Human PINK1-WT were cloned into the pBad/HisB vector that already encoded for the CyPet/YPet FRET-pair. The vector was transformed into TOP10 chemically competent *E. coli* cells (ThermoFisher, USA). Transformed cells were grown overnight at 37°C on LB agar plates containing 100 μg/mL ampicillin. One transformant colony was selected and grown overnight at 37°C in 120 mL of LB medium containing 100 μg/mL ampicillin. A total of 6 L LB media was sub-inoculated with 20 mL of overnight culture and grown to an OD_600_ of 0.7 at 37°C. Cultures were induced by addition of 0.02% (v/v) L-arabinose for 8 h at 24°C. After induction, cells were harvested by centrifugation in a Beckman JLA8.1000 rotor (6,900 × *g*, 20 min, 4°C), flash frozen in liquid nitrogen, and stored at −80°C. Harvested cells were thawed on ice and resuspended in a 4:1 buffer volume to cell pellet weight ratio in resuspension buffer (50 mM Tris-HCl pH 8.0, 500 mM NaCl, 20% glycerol, 10 μg/mL DNase, 1 mM PMSF, two EDTA-free protease inhibitor cocktail tablets). Resuspended cells were lysed using an Emulsiflex with a maximum pressure of 40 kPSI. Following cell lysis, the lysate was subjected to centrifugation using a Beckman TI45 rotor (31,300 × *g*, 20 min, 4°C) to pellet cell debris and unlysed cells. The supernatant was incubated with 1% (v/v) Triton X-100 at 4°C for 30 min with stirring. Supernatant was then passed through 1 mL settled HisPur™ cobalt resin (ThermoFisher, USA) by gravity flow to allow binding of FRET- PINK1-His to the resin. Protein was eluted (50 mM Tris-HCl pH 8.0, 500 mM NaCl, 20% glycerol, 250 mM imidazole), pooled, and concentrated for loading onto the Superdex 200 column for size exclusion chromatography. Size exclusion chromatography fractions were analyzed by SDS-PAGE and fractions containing FRET-PINK1 protein were pooled and concentrated. Concentrated sample was aliquoted, flash frozen with liquid nitrogen, and purified HsFRET-PINK1(70-134) was stored at −80 °C for subsequent use.

### PINK1 TM expression and purification

The sequence of PINK1 TM domain (amino acids 89-111) was codon optimized for *E. coli* expression and cloned into pMAL-c2 vector (New England Biolab) with N-terminal Maltose Binding Protein (MBP) followed by a tobacco etch virus (TEV) cleavage site. The vector was transformed into DH5α cells and the protein was induced with 0.5 mM IPTG and expressed for 3 days at 24°C. Cells were harvested, resuspended in 20 mM KPO_4_ (pH 8), 120 mM NaCl, 50 mM glycerol, 1 mM EDTA, 1 mM PMSF, 1 mM DTT and lysed using an Emulsiflex with a maximum pressure of 40 kPSI. 0.5% Triton X-100 was added post-lysis and cell debris were removed by centrifugation at 40,000 g for 30 min at 4 °C. The supernatant was loaded onto amylose resin (Amylose Resin High Flow, NEB), equilibrated with 20 mM KPO_4_, pH 8.0, 120 mM NaCl, 1 mM EDTA buffer and the protein was eluted with 40 mM maltose in equilibration buffer. MBP tag was cleaved off by MBP-PINK1 incubation with recombinant TEV protease (1.5 mg of TEV per 30 mg of fusion protein) at 16 °C for 4-8 days. To extract PINK1 TM segment 1/6 of the sample volume of 60% w/v trichloroacetic acid (TCA) was added to protein mixture and incubated for 30 min on ice. The precipitate was pelleted for 10 min at 10,000 × *g*, rinsed 3 times with ddH_2_O, resuspended in 50:50 isopropanol:chloroform and mixed with a homogenizer. To this mixture, 1-2 mL of ddH_2_O was added into each tube and incubated overnight allowing for separation of the organic and aqueous layers. The organic layer was transferred into a clean tube and fresh 1-2 mL of was aliquoted into a sample and left overnight at room temperature. This separation was repeated until all white precipitate was removed and organic phase was considered clean. Organic layers were combined and dried down under nitrogen or argon gas. The PINK1 peptide was resuspended in ~6-8 mL of 7 M guanidine-HCl, 50 mM KPO_4_ buffer (pH 8) and injected onto an Agilant Zorbax SB-300 C8 silica based, stainless steel 25 cm × 1 cm column, which was preheated to 60 °C. The column ran at 60 °C with a flow rate of 1 mL/min. An isopropanol gradient (20% – 80%) against 0.05% TFA/water was used to elute the protein. PINK1 TM typically eluted at ~50% isopropanol. Determination of fractions containing the peptide was established by running 6% urea gels, which were visualized through silver staining.

### MBP-PGAM5 expression and purification

The sequence of the PGAM5 TM region (amino acids 1-46) was cloned into *E. coli* expression vector pET-25b(+) (Novagen) with N-terminal MBP and C-terminal Thioredoxin 1 followed by a triple FLAG-tag and a C-terminal hexahistidine-tag. The vector was transformed into chemical competent Rosetta™ 2 (DE3) cells (Novagen), grown in LB medium. Expression of the protein was induced with 0.3 mM IPTG and expressed for 2 hours at 37 °C. Cells were harvested by centrifugation at 3500 rpm for 15 min at 4°C and resuspended in 20 mM HEPES pH 7.4, 150 mM NaCl, 5 mM MgCl_2_, 10 % glycerol, 1 mM PMSF, 5 mM β-mercaptoethanol. Prior to lysis, 200 μg/mL lysozyme, 1 mM PMSF and benzonase (2.5 ku, Merck Millipore) were added and cells were lysed using Emulsiflex (Avestin) with a maximum pressure of 15 kPSI (100 MPa). Crude membranes were obtained by ultracentrifugation at 29,000 rpm for 45 min at 4°C. The membrane pellet was resuspended in 50 mM HEPES pH 7.4, 150 mM NaCl, 5 mM MgCl_2_, 10% glycerol, 1 mM PMSF, 5 mM β-mercaptoethanol. MBP-PGAM5 was solubilized from the crude membranes with 1.5 % DDM for 1 hour on a rotating wheel at room temperature. Extraction of MBP-PGAM5 from membrane debris was done by ultracentrifugation at 29,000 rpm for 1 hour at 4°C. Cleared extract was batch incubated with Ni-NTA beads (Macherey-Nagel) for 1 hour on a rotating wheel at room temperature for His-tag affinity purification. Bound MBP-PGAM5 was washed with 50 mM HEPES pH 7.4, 300 mM NaCl, 10% glycerol, 50 mM imidazole, 0.05 % DDM and eluted with 50 mM HEPES pH 7.4, 300 mM NaCl, 10% glycerol, 400 mM imidazole, 0.05 % DDM. Determination of fractions containing the peptide was established by SDS-PAGE running 12% acrylamide gels, which were visualized through Coomassie staining.

### MBP-PGAM5 cleavage assay

5 μg (4 μM) of *E. coli* purified MBP-PGAM5 was incubated with either 0.44 μg (0.7 μM) PARL or 0.44 μg (0.7 μM) catalytic inactive PARL-S277A purified from *P. pastoris* for 1.5 hours at 30°C in cleavage buffer containing 50 mM Tris pH 8.0, 150 mM NaCl, 10 % glycerol, 0.3 % DDM. Determination of peptide cleavage was established by SDS-PAGE using 12% acrylamide gels, which were visualized through Coomassie staining.

### N-terminal sequencing by Edman degradation

8-16 μg of *E. coli* purified MBP-PGAM5 was incubated with 0.4 μg of *P. pastoris* purified PARL for 2 hours at 37°C in cleavage buffer containing 50 mM Tris pH 8.0, 150 mM NaCl, 10 % glycerol, 0.3 % DDM. Protein fragments were separated by SDS-PAGE running 12% acrylamide gels and transferred to a PVDF membrane by wet blot (glycine buffer) for 1 hour at 100 V. Protein fragments were stained with Coomassie overnight and the C-terminal fragment (CTF) was then analyzed in 4 cycles by Edman degradation (TOPLAB, Germany).

### FRET-based protease kinetic assay

Assays with FRET-PINK1^70-134^ were conducted as previously described (Arutyunova et al, 2014). For EDANS/Dabcyl 10-mer IQ peptides (PINK1, PGAM5, Smac), lyophilized peptides were initially dissolved in DMSO to obtain a stock solution. The IQ peptide substrates in a concentration range of 0.1 to 70 μM were incubated with activity assay buffer (50 mM Tris-HCl pH 7.0, 150 mM NaCl, 10 % glycerol, 0.1% DDM) in a 384-well black-bottomed plate at 37°C for 30 min in a multi-well plate reader (SynergyMx, BioTek). For all concentrations of IQ peptide, the DMSO was kept constant at 5%. Following pre-incubation, PARL was added to a final concentration of 0.8 μM to initiate the cleavage reaction. Fluorescence readings were taken every 3 min over a 3 h time course at ƛ_ex_ = 336 nm and ƛ_em_ = 490 nm. The initial velocity was determined from the fluorescence readings over the time course. For each substrate concentration, a no-enzyme control was subtracted to eliminate background fluorescence changes not related to substrate cleavage. Relative fluorescence units were converted to concentration (μM) by determining the maximum change in fluorescence observed for each substrate concentration when fully digested. GraphPad Prism software was used for Michaelis-Menten analysis of kinetic curves. Minimum of three experimental replicates with two technical replicates used for data analysis.

### Reconstitution in proteoliposomes

*E. coli* polar lipids (Avanti), 400 μg in chloroform, were dried under nitrogen stream in a glass tube to yield a thin film of lipid. The tube was incubated overnight in a desiccator to completely remove all traces of solvent. 50 μl of water and DDM detergent was added to the lipid film for resuspension at room temperature for 10 min, followed by the addition of purified PARL (400 μg) in 50 mM Tris-HCl, pH 8.0, 150 mM NaCl, 20 % glycerol, 0.1 % DDM, to yield the final weight ratio of 1 PARL:1 lipid:2 detergent. The detergent was slowly removed by the addition of SM2 Biobeads (Bio-Rad, USA) while stirring on ice for 6 hours to allow for the generation of proteoliposomes (PL); the process was controlled by the regimen of Biobead addition. To purify the PL, 50%:20% sucrose density gradient ultracentrifugation was used. A TAMRA probe (ThermoFisher) was used to determine the orientation of reconstituted PARL, which specifically and covalently labels serine residues of an enzymatically active serine protease. 2 μM of TAMRA probe was added to PL samples and incubated for 1 hour. The same amount of PL, but with 1% DDM added to dissolve lipid vesicles was used as a benchmark for 100 % accessible protein amount. The reaction was quenched with addition of SDS-containing sample buffer and the protein samples were visualized with SDS-PAGE followed by fluorescent gel scanning and Coomassie staining. Coomassie staining was used to normalize the amount of protein loaded.

### Activity assay in proteoliposomes

The activity assay with PARL reconstituted in PL was performed the same way as for PARL in DDM with the only difference being the activity buffer, where DDM was omitted (50 mM Tris-HCl pH 7.0, 150 mM NaCl, 10 % glycerol). For inhibitory studies, PARL in PL (0.8 μM) was incubated with inhibitors (20 μM) in activity buffer for 30 min and then the proteolytic reaction was started with the addition of substrate (5 μM).

### Molecular modeling

Human PARL was modelled using iTASSER with PARL isoform 1 as the template, without any additional restraints(Yang et al, 2015). Residues 1-167 were removed from the modeling due to low homology with bacterial rhomboid protease crystal structures.

### Multiplex substrate profiling by mass spectrometry

Multiplex substrate profiling by mass spectrometry (MSP-MS) assays were performed in quadruplicate. 1 μM of PARLΔ77 was incubated with an equimolar mixture of 228 synthetic tetradecapeptides at a final concentration of 0.5 μM for each peptide in 50 mM Tris-HCl pH 7.0, 150 mM NaCl, 10% glycerol, with or without 0.1% DDM. For each assay, 20 μL of the reaction mixture was removed after 0, 60 and 240 minutes of incubation. Enzyme activity was quenched by adding GuHCl (MP Biomedicals) to a final concentration of 6.4 M and samples were immediately stored at −80 °C. All samples were desalted using C18 spin columns and dried by vacuum centrifugation.

Approximately 2 μg of peptides was injected into a Q-Exactive Mass Spectrometer (Thermo) equipped with an Ultimate 3000 HPLC. Peptides were separated by reverse phase chromatography on a C18 column (1.7 μm bead size, 75 μm × 25 cm, 65 °C) at a flow rate of 300 nL/min using a 60-minute linear gradient from 5% to 30% B, with solvent A: 0.1% formic acid in water and solvent B: 0.1% formic acid in acetonitrile. Survey scans were recorded over a 150–2000 m/z range (70,000 resolutions at 200 m/z, AGC target 3×10^6^, 100 ms maximum). MS/MS was performed in data-dependent acquisition mode with HCD fragmentation (28 normalized collision energy) on the 12 most intense precursor ions (17,500 resolutions at 200 m/z, AGC target 1×10^5^, 50 ms maximum, dynamic exclusion 20 s).

Data was processed using PEAKS 8.5 (Bioinformatics Solutions Inc.). MS^2^ data were searched against the 228 tetradecapeptide library sequences with decoy sequences in reverse order. A precursor tolerance of 20 ppm and 0.01 Da for MS^2^ fragments was defined. No protease digestion was specified. Data were filtered to 1% peptide level false discovery rates with the target-decoy strategy. Peptides were quantified with label free quantification and data are normalized by medians and filtered by 0.3 peptide quality. Missing and zero values are imputed with random normally distributed numbers in the range of the average of smallest 5% of the data ± SD. IceLogo software was used for visualization of amino-acid frequency surrounding the cleavage sites using the sequences of all cleavage products whose fold change > 8 and qvalue < 0.05 (by Student t-test) in 60 min. Amino acids that were most frequently observed (above axis) and least frequently observed (below axis) from P4 to P4ʹ positions were illustrated. Norleucine (Nle) was represented as ‘n’ in the reported profiles. Mass spectrometry data and searching results have been deposited in MassIVE with accession number, MSV000085295.

## Acknowledgements

We thank Baptiste Cordier for advice and technical support. Funding: M.J.L. gratefully acknowledges support from the Parkinson Society of Canada, and by grants from the Canadian Institutes of Health Research (MOP-93557, Funder Id:10.13039/501100000030), Natural Sciences and Engineering Research Council of Canada – NSERC (RGPIN-2016-06478), Neuroscience and Mental Health Institute (Johnston Family Endowment Fund, and the Brad Mates Foundation), Parkinson Alberta, and the Department of Biochemistry, University of Alberta. M.J.L. was supported in part by funding from Alberta Innovates Health Solutions and the Parkinson’s Society of Canada New Investigator Program. M.K.L. was supported by the grant Le2749/1-2 of the Deutsche Forschungsgemeinschaft (German Research Foundation) as part of FOR2290.

## Author contributions

Experiments were devised by JL, EA, AO, ML. LL and EA conducted the kinetic assays, MM, LL, ET, EA, and RB made PARL protease and PINK1 protein. EA conducted proteoliposome assays, VS conducted MBP-PGAM5 cleavage assay and Edman degradation assay, ZJ conducted substrate specificity profiling. HSY conducted EM analysis of liposomes. The manuscript was written by JL and edited by all authors.

## Supplemental information

**Supplemental Figure 1.**
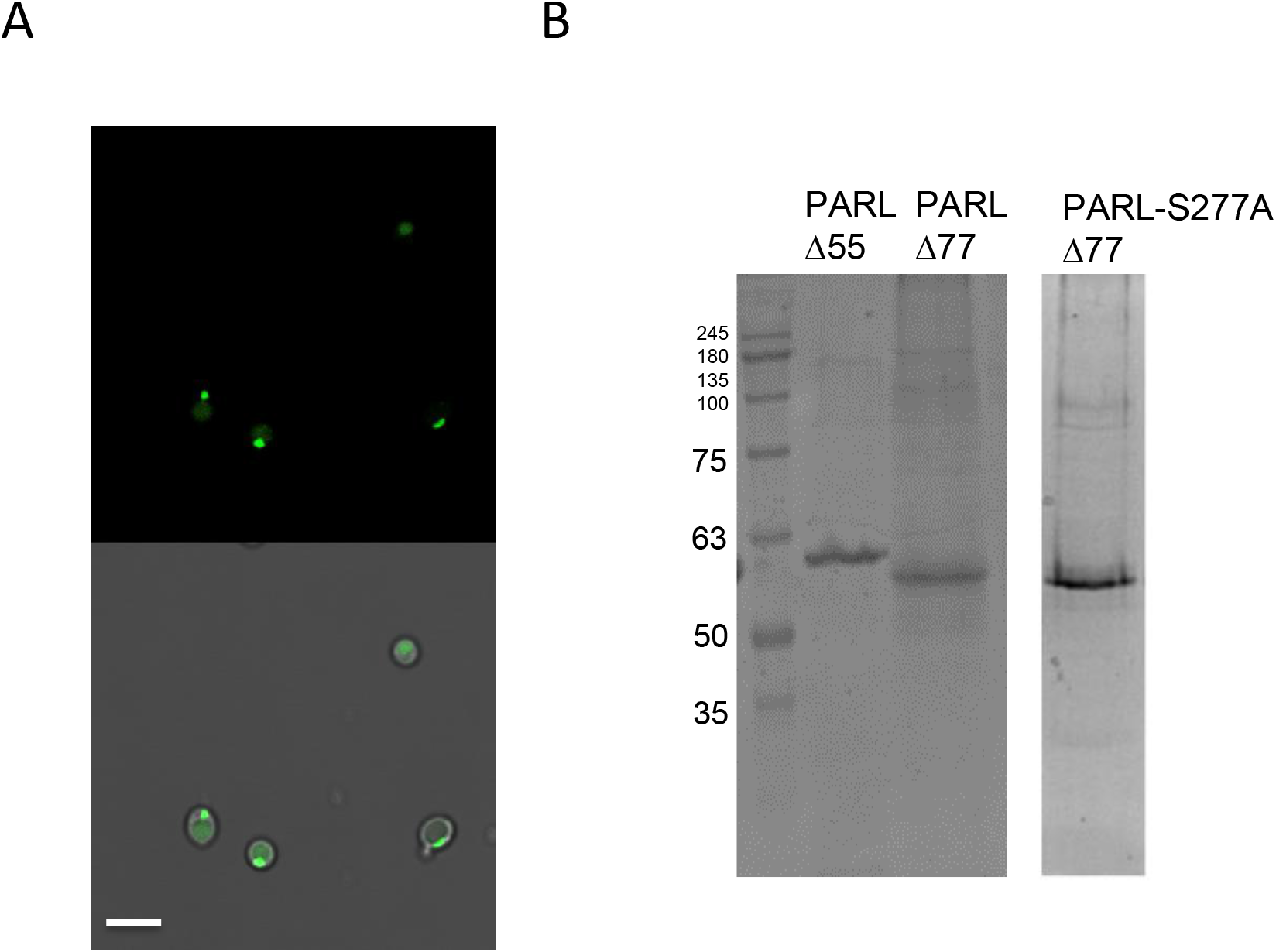
**A.** PARLΔ77-GFP in *Pichia* showing the enzyme localizes to the vacuole compartment. The PARL (green) direct fluorescence signal (Top) was merged with a phase-contrast image of the cells (gray). Images were collected on a Zeiss LSM 410 confocal microscope. The scale bar represents 4 μm. **B.** Recombinant human PARL was expressed in *P. pastoris* either truncated at residue Δ55 or Δ77. An inactive PARL-Δ77-S277A mutation was also generated.

**Supplemental Figure 2.**
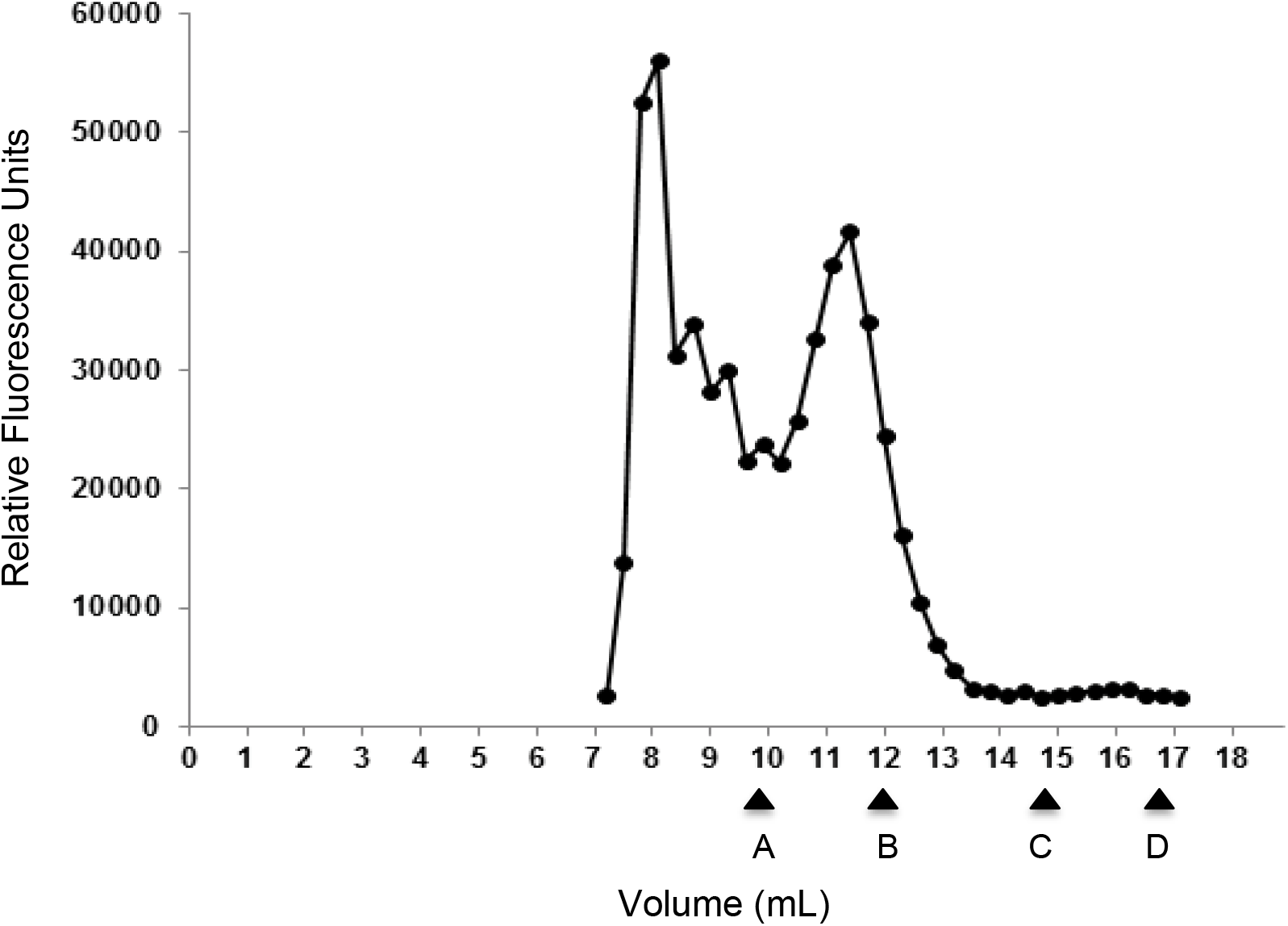
FSEC of PARL-GFP in n-dodecyl β-D-maltopyranoside (DDM) reveals a homogenous sample. Filter-centrifuged crude membranes of PARL-GFP (100 μL), solubilized in DDM were loaded on a Superdex 200 (10×300) column in 50 mM KPO4, 5% glycerol, 0.1 M NaCl, 10 mM βME, 0.1% DDM.. Fractions were collected every 300 μL. Relative fluorescence was measured separately in a fluorimeter. The column void volume is 8.32 ml. The arrows represent Molecular Weight Standards: A-Thyroglobulin (MW, 670 kDa; Stokes radius 85 Å); B – γ-globulin (MW 158 kDa; Stokes radius 52.9 Å); C – Ovalbumin (MW 44 kDa; Stokes radius 30.5 Å); D – myoglobin (MW 17 kDa, Stokes radius 20.7 Å).

**Supplemental Figure 3.**
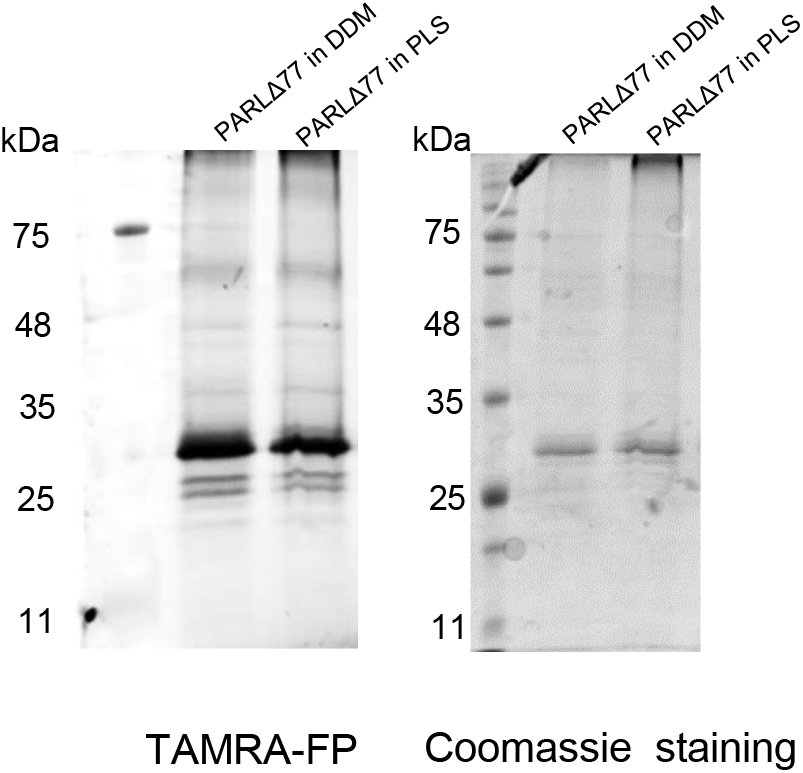
SDS-PAGE of PARL with TAMRA probe. To determine orientation of PARL in the proteoliposomes, PARL in PLs was incubated with TAMRA and then separated on a 14% SDS-PAGE gel. The fluorescent TAMRA probe was visualized using an imager (left) and also stained with Coomassie blue (right).

**Supplemental Figure 4.**
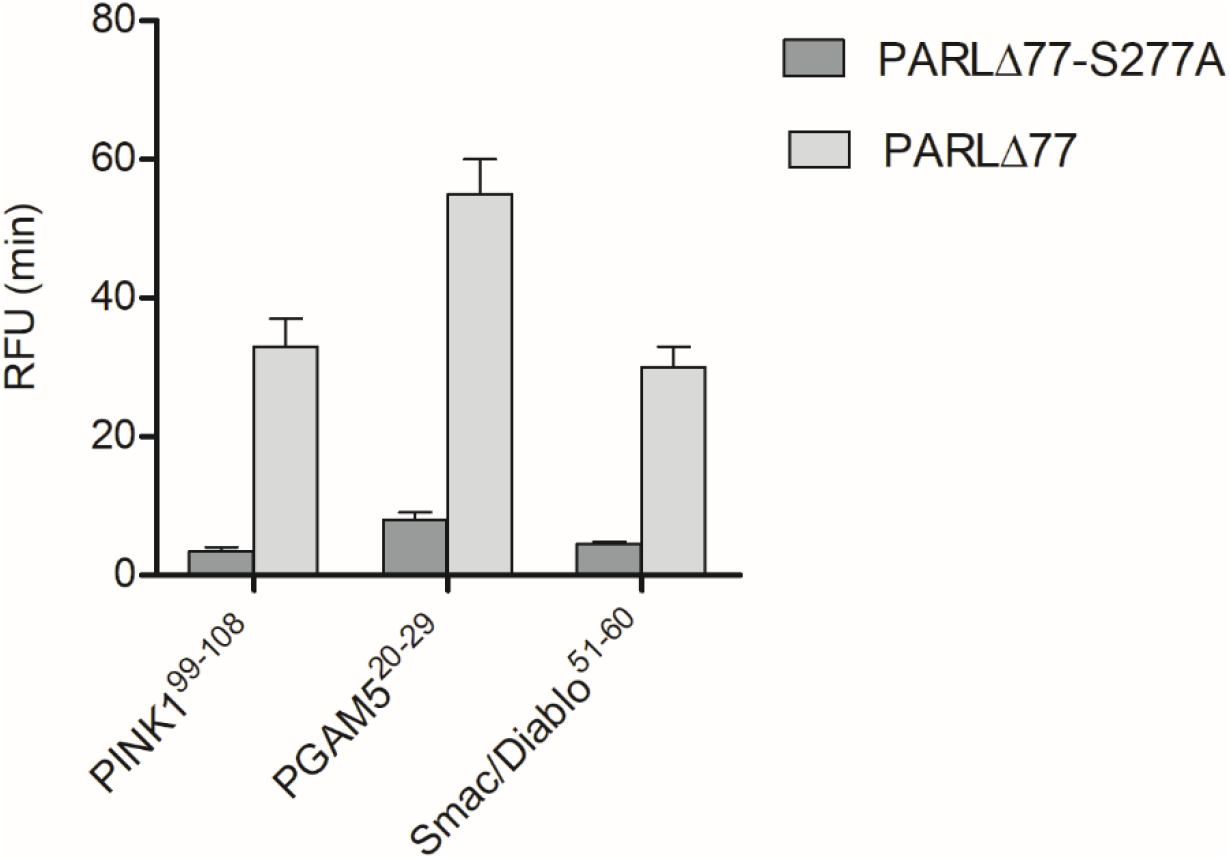
Inactive PARLΔ77-S277A in proteoliposomes shows negligible activity with PARL substrates. The activity of PARLΔ77 and PARLΔ77-S277A was measured with 2.3μM of each substrate.

**Supplemental Figure 5.**
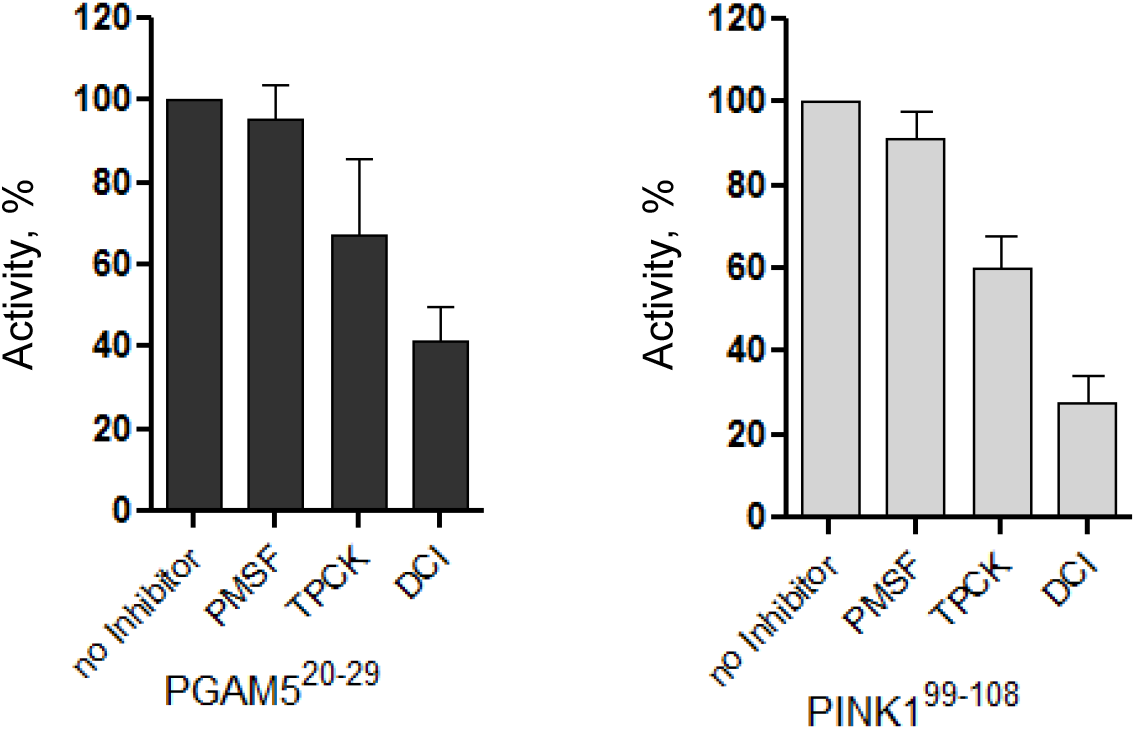
PARL is not inhibited by commercially available serine protease inhibitors. Bar graphs for percent activity of proteoliposome reconstituted PARLΔ77 with 20 μM of inhibitor added prior to the addition of IQ peptide substrates PINK1 and PGAM5. Experiments were conducted in duplicate with an N=3. Values are represented as mean ±SEM. * p < 0.05, ** p < 0.005, *** p < 0.0005, n.s. denotes no significance.

**Supplemental Table 1.** Catalytic parameters of PARLΔ55 and PARLΔ77 with FRET-PINK1^70-134^ substrate.

|  | PARLΔ55 | PARLΔ77 |
| --- | --- | --- |
| $K_M$ (μM) | 3±1 | 5±1 |
| $k_{cat}$ (h<sup>-1</sup>) | 0.46±0.09 | 0.43±0.06 |
| $k_{cat}/K_M$ (μM<sup>-1</sup>h<sup>-1</sup>) | 0.16±0.09 | 0.09±0.03 |

**Supplemental Table 2.**
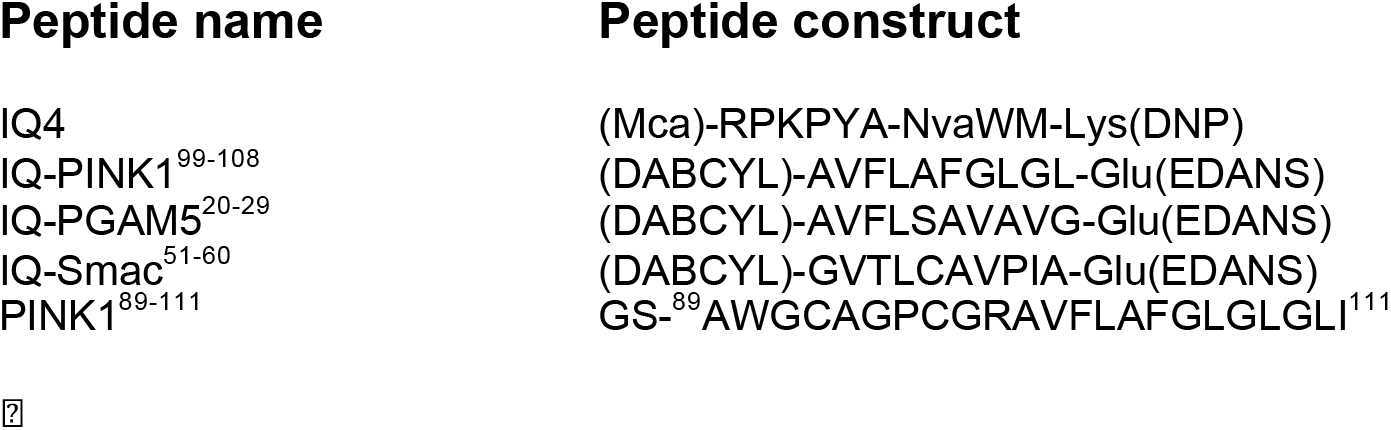
Synthetic peptides used in this study

**Supplemental Table 3.** Catalytic parameters of FL PARLΔ55-mediated cleavage of IQ-peptide substrates. Experiments were conducted in duplicate with an N=4-5. Values are represented as mean ±SEM.

| PARLΔ55<br>(Full length ) | PINK1<sup>99-108</sup> | PGAM5<sup>20-29</sup> | Smac/Diablo<sup>51-60</sup> |
| --- | --- | --- | --- |
|  | n = 4 | n = 5 | n = 4 |
| $K_M$ (μM) | 1.9 ± 0.4 | 0.47 ± 0.07 | 6 ± 1 |
| $k_{cat}$ ($h^{-1}$) | 0.42 ± 0.03 | 0.46 ± 0.02 | 1.3 ± 0.1 |
| $k_{cat}/K_M$ (μM<sup>-1</sup>h<sup>-1</sup>) | 0.22 ± 0.08 | 1.0 ± 0.3 | 0.3 ± 0.1 |

**Supplemental Table 4.** Catalytic parameters of β-cleavage PARLΔ77-mediated cleavage of IQ-peptide substrates. Experiments were conducted in duplicate with an N=4-5. Values are represented as mean ±SEM.

| PARLΔ77<br>(β-cleavage) | PINK1<sup>99-108</sup> | PGAM5<sup>20-29</sup> | Smac/Diablo<sup>51-60</sup> |
| --- | --- | --- | --- |
|  | n = 4 | n = 5 | n = 4 |
| $K_M$ (μM) | 3.3 ± 0.5 | 1.6 ± 0.2 | 5 ± 1 |
| $k_{cat}$ ($h^{-1}$) | 1.3 ± 0.1 | 0.98 ± 0.04 | 1.2 ± 0.1 |
| $k_{cat}/K_M$ (μM<sup>-1</sup>h<sup>-1</sup>) | 0.4 ± 0.1 | 0.6 ± 0.1 | 0.3 ± 0.1 |

**Supplemental Table 5.**
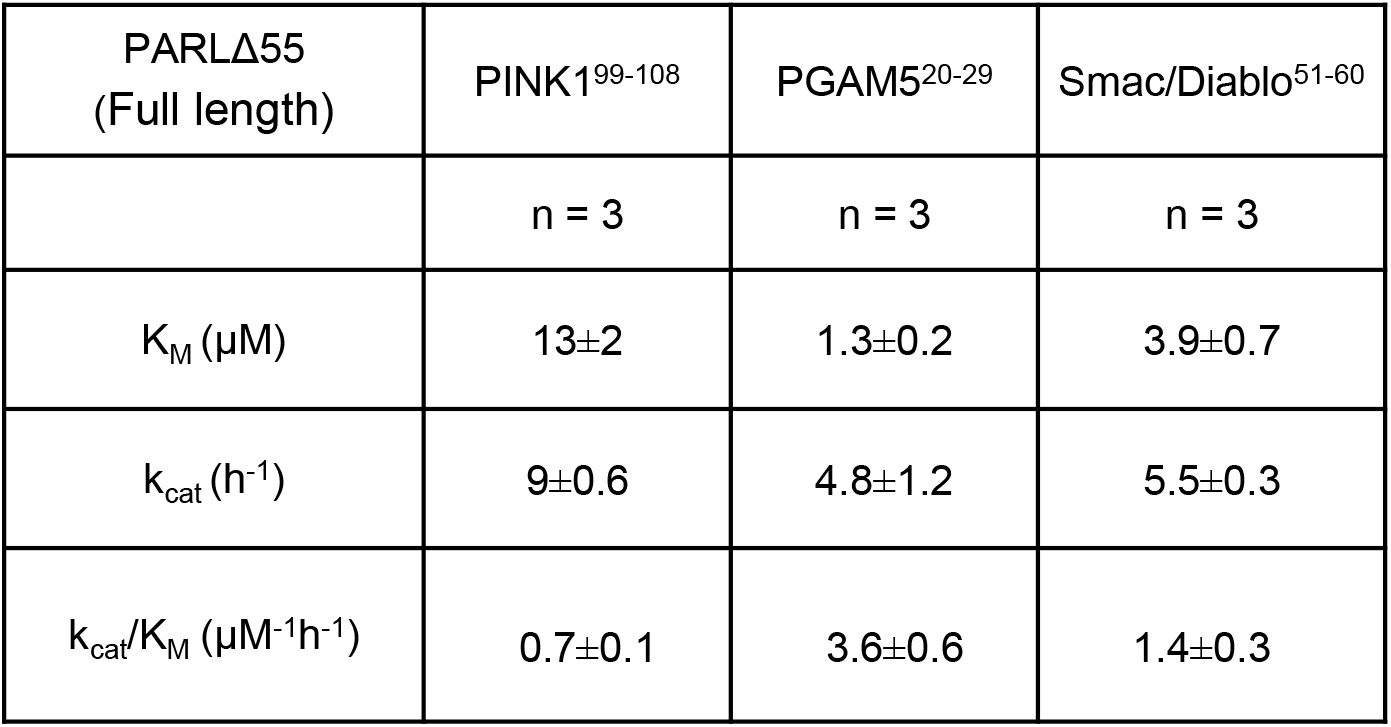
Specific proteolytic activity of full length PARLΔ55 in proteoliposomes (POPE/POPC/CL). Experiments were conducted in duplicate with an N=3. Values are represented as mean ±SEM.

| PARLΔ55<br>(Full length) | PINK1<sup>99-108</sup> | PGAM5<sup>20-29</sup> | Smac/Diablo<sup>51-60</sup> |
| --- | --- | --- | --- |
|  | n = 3 | n = 3 | n = 3 |
| $K_M$ (μM) | 13±2 | 1.3±0.2 | 3.9±0.7 |
| $k_{cat}$ ($h^{-1}$) | 9±0.6 | 4.8±1.2 | 5.5±0.3 |
| $k_{cat}/K_M$ (μM<sup>-1</sup>$h^{-1}$) | 0.7±0.1 | 3.6±0.6 | 1.4±0.3 |

**Supplemental Table 6.** Specific proteolytic activity of β-cleaved PARLΔ77 in proteoliposomes (POPE/POPC/CL). Experiments were conducted in duplicate with an N=3. Values are represented as mean ±SEM.

| PARLΔ77<br>(β-cleavage) | PINK1<sup>99-108</sup> | PGAM5<sup>20-29</sup> | Smac/Diablo<sup>51-60</sup> |
| --- | --- | --- | --- |
|  | n = 4 | n = 5 | n = 4 |
| $K_M$ (μM) | 8±1 | 0.8±0.2 | 17±4 |
| $k_{cat}$ ($h^{-1}$) | 28±2 | 21±1 | 58±7 |
| $k_{cat}/K_M$ (μM<sup>-1</sup>$h^{-1}$) | 3.5±0.6 | 26±8 | 3.4±1 |

**Supplemental Table 7.** Specific proteolytic activity of PARLΔ77 protease in POPE/POPC proteoliposomes (lacking CL). Experiments were conducted in duplicate with an N=3. Values are represented as mean ±SEM.

| PARLΔ77<br>(β-cleavage) | PINK1<sup>99-108</sup> | PGAM5<sup>20-29</sup> | Smac/Diablo<sup>51-60</sup> |
| --- | --- | --- | --- |
|  | n=3 | n=3 | n=3 |
| $K_M$ (μM) | 14±5 | 8±3 | 3.8±1 |
| $k_{cat}$ ($h^{-1}$) | 7±1 | 9±2 | 4.8±0.3 |
| $k_{cat}/K_M$ (μM<sup>-1</sup>h<sup>-1</sup>) | 0.5±0.2 | 1.0±0.3 | 1.2±0.3 |

**Supplemental Table 8.** The fold change in turnover rate and catalytic efficiency is shown between β-cleavage (PARLΔ77) PARL and full length (PARLΔ55) either in **A.** detergent and **B.** reconstituted in proteoliposomes.

**A**
| Fold change<br>(PARL det)<br>PARLΔ77/PARLΔ55 | $K_{cat}$ | $K_m/K_{cat}$ |
| --- | --- | --- |
| PINK1 | 3 | 2 |
| PGAM5 | 2 | 0.6 |
| Smac | 1 | 1 |

| Fold change<br>(PARL in PL)<br>PARLΔ77/PARLΔ55 | $K_{cat}$ | $K_m/K_{cat}$ |
| --- | --- | --- |
| PINK1 | 3 | 5 |
| PGAM5 | 4 | 7 |
| Smac | 10 | 2 |

